# Structure of a Membrane Tethering Complex Incorporating Multiple SNAREs

**DOI:** 10.1101/2023.01.30.526244

**Authors:** Kevin A. DAmico, Abigail E. Stanton, Jaden D. Shirkey, Sophie M. Travis, Philip D. Jeffrey, Frederick M. Hughson

**Affiliations:** Department of Molecular Biology, Princeton University, Princeton, NJ 08544

## Abstract

Most membrane fusion reactions in eukaryotic cells are mediated by membrane tethering complexes (MTCs) and SNARE proteins. MTCs are much larger than SNAREs and are thought to mediate the initial attachment of two membranes. Complementary SNAREs then form membrane-bridging complexes whose assembly draws the membranes together for fusion. Here, we present a cryo-EM structure of the simplest known MTC, the 255-kDa Dsl1 complex, bound to the two SNAREs that anchor it to the endoplasmic reticulum. N-terminal domains of the SNAREs form an integral part of the structure, stabilizing a Dsl1 complex configuration with remarkable and unexpected similarities to the 850-kDa exocyst MTC. The structure of the SNARE-anchored Dsl1 complex and its comparison with exocyst reveal what are likely to be common principles underlying MTC function. Our structure also implies that tethers and SNAREs can work together as a single integrated machine.

## INTRODUCTION

Cargo in eukaryotic cells is transported between organelles, and to and from the plasma membrane, in vesicles and other membrane-bound carriers. The initial contact between vesicle and target membranes is mediated by two classes of organelle-specific tethering factors: extended coiled coil homodimers (e.g., golgins) and multisubunit tethering complexes (MTCs)^1,2^. Both classes of tether link membranes by binding to determinants such as lipids, small GTPases, SNAREs, and vesicle coat proteins. MTCs also carry out functions that transcend simple tethering but are crucial for membrane fusion, such as chaperoning SNARE assembly and promoting membrane curvature^3,4^. MTCs may thereby orchestrate the entire cargo delivery process, from membrane recognition through to SNARE-mediated membrane fusion.

The largest family of MTCs is the CATCHR (complexes associated with tethering containing helical rods) family, whose members function in anterograde and retrograde trafficking throughout the secretory system^5,6^. A second family, HOPS/CORVET, mediates the tethering of endolysosomal membranes^7^. The CATCHR family MTCs are composed of 3-8 different subunits: the Dsl1 complex has 3, EARP and GARP each have 4, and COG and exocyst each have 8. Nearly all of these subunits are structurally homologous, with a roughly 650-residue C-terminal region consisting of a rod-like series of helical bundle domains denoted A-E^8–16^. This CATCHR fold is also found in the monomeric tethering proteins Munc13, which tethers synaptic vesicles to the pre-synaptic membrane, and myosin V, which tethers membrane cargo to the cytoskeleton^17,18^.

The assembly of CATCHR-family MTCs depends on N-terminal sequences that, through antiparallel coiled-coil interactions, generate subunit pairs^5,10,14,19^. Indeed, a landmark 4.4-Å cryo-EM structure of the 850-kDa hetero-octameric exocyst complex from *Saccharomyces cerevisiae*^11^ revealed 4 such pairs, further organized into two 4-subunit subassemblies. These subassemblies interact, largely via their CATCHR domains, to generate an elaborate architecture with overall dimensions of approximately 13 x 32 nm. In addition to their structural roles, the CATCHR domains of exocyst have been implicated in a wide array of intermolecular interactions with partners including the phospholipid PI(4,5)P_2_, the Rab GTPase Sec4, multiple Rho GTPases, and multiple SNAREs^20^. A pleckstrin homology domain, not observed in the cryo-EM structure, contains additional partner-binding sites. Nonetheless, despite remarkable progress, the large size, complex architecture, and broad interactome of exocyst have made it challenging to elucidate how it mediates membrane tethering, SNARE assembly, and fusion.

All three of these core functions are also supported by the much-simpler Dsl1 complex^21,22^. This is particularly interesting since the Dsl1 complex would appear to be the least exocyst-like of the CATCHR-family MTCs. At 255 kDa, it is 70% smaller than exocyst, and it consists of just 3 subunits: Dsl1, Tip20, and Sec39. Moreover, only the Dsl1 and Tip20 subunits possess the canonical N-terminal coiled coil and C-terminal CATCHR domains; Sec39 is instead a rod-like α-solenoid^14,23^. All 3 subunits are encoded by essential genes and are required for retrograde trafficking from the Golgi to the endoplasmic reticulum (ER)^24–27^.The ability of the Dsl1 complex to tether Golgi-derived COPI-coated vesicles to the ER is consistent with its known interactome, which includes two subunits of the COPI coat (α-COP and δ-COP) and two ER-anchored SNARE proteins (Sec20 and Use1)^23,28,29^.

SNAREs are much smaller than MTCs, and most of them share a canonical structure consisting of a structured N-terminal domain (NTD), a SNARE motif, and a C-terminal transmembrane anchor^30^. SNARE motifs are roughly 60 residues in length and are unstructured in isolation^31^. The formation of a fusogenic SNARE complex entails the coupled folding and assembly of 4 SNARE motifs to form a stable, membrane-bridging 4-helix bundle^32,33^. The 4 complementary SNARE motifs – one each from the R, Qa, Qb, and Qc families – are generally present in 4 different SNARE proteins, although a few SNARE proteins such as SNAP-25 and its yeast homolog Sec9 contain both Qb and Qc SNARE motifs^31,34^. In all cases, the assembling SNARE motifs exert force to pull the two membranes together^35^. The NTDs, by contrast, play more indirect roles in membrane fusion. One role, important for the regulation of SNARE assembly, is to interact intramolecularly with the SNARE motif^36,37^. A second role is to interact intermolecularly with other components of the membrane fusion machinery, including MTCs^23,38^. Indeed, ER SNAREs bind to the Dsl1 complex by means of their NTDs, not their SNARE motifs^23,39^. Accordingly, they might assemble into membrane-bridging complexes, mediate fusion, and disassemble again, all while remaining bound to the Dsl1 complex.

The 3 subunits of the Dsl1 complex and the 2 SNAREs Sec20 (a Qb-SNARE) and Use1 (a Qc-SNARE) combine to form stable hetero-pentamers that can be co-immunoprecipitated from yeast lysates and likely represent the Dsl1 complex in its ER-anchored state^26^. Here, we have reconstituted this complex, lacking only the transmembrane anchors of the two SNAREs and a non-essential C-terminal luminal domain of Sec20, and have determined its structure using single-particle cryo-EM. This is to our knowledge the first structure of an intact MTC bound to SNAREs or, indeed, to any other proteins. The SNAREs play a key structural role, interacting via their NTDs to form a tether:SNARE complex with a pronounced resemblance to exocyst. Compromising the assembly of this complex is lethal in yeast, suggesting that the 3 Dsl1 subunits and 2 SNAREs function in intimate collaboration to help orchestrate vesicle capture, SNARE assembly, and membrane fusion.

## RESULTS

### Structure of the Dsl1:Qb:Qc complex

The three subunits of the *S. cerevisiae* Dsl1 complex (Dsl1, Tip20, and Sec39) were coexpressed in bacteria with the cytoplasmic portions of the ER Qb- and Qc-SNAREs (Sec20 and Use1) (Fig. 1a). The resulting complex, hereafter called Dsl1:Qb:Qc, was stable and monodisperse (Fig. 1b). Earlier negative-stain studies of the Dsl1 complex in the absence of SNAREs suggested that it contains flexible hinges and adopts a range of conformations^23^. By contrast, negative stain EM examination of Dsl1:Qb:Qc revealed a much more uniform conformation (Fig. 1c). Particles were triangular in shape with a maximum dimension of approximately 25 nm.

**Fig. 1:**
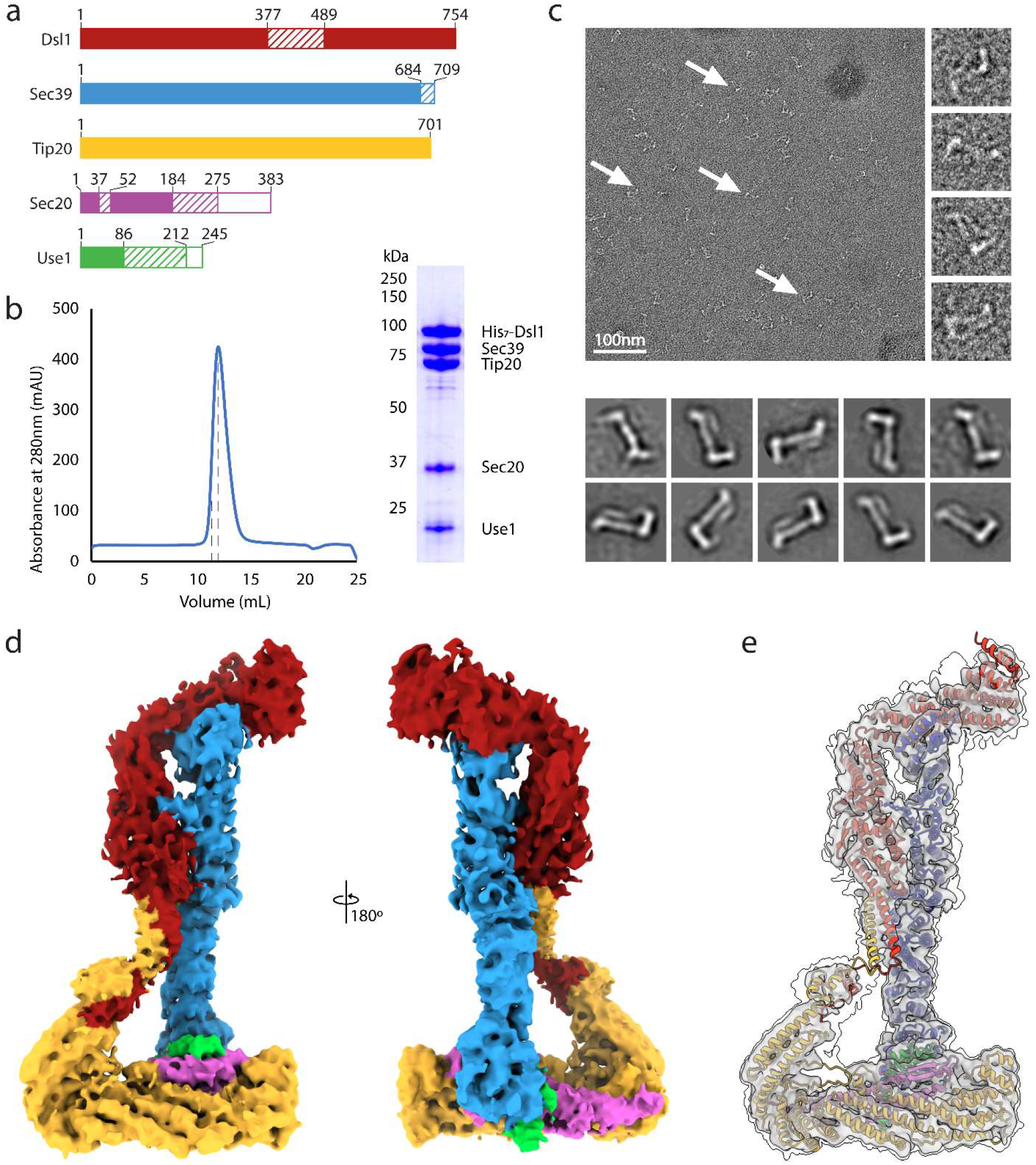
The Dsl1 complex bound to Sec20 and Use1. **a**, Schematic of the *S. cerevisiae* polypeptides used in this work: the Dsl1 complex subunits Dsl1, Sec39, and Tip20, the Qb-SNARE Sec20, and the Qc-SNARE Use1. Solid regions are modeled in the Dsl1:Qb:Qc structure, striped regions are disordered, and open regions were not included in the protein expression plasmids. Color coding is consistent throughout. **b,** Size-exclusion chromatography (left) of the Dsl1:Qb:Qc complex. Dashed lines indicate the fraction visualized (right) on a 10% polyacrylamide SDS-PAGE gel. **c,** Negative-stain EM of the Dsl1:Qb:Qc complex. Shown are a representative field (top) and 2D averages (bottom). **d,** Composite of locally-refined cryo-EM density maps at 4.5 Å resolution. e, Overlay of the composite map at two contour levels and Dsl1:Qb:Qc model.

To determine a higher resolution structure of Dsl1:Qb:Qc, we used single-particle cryo-EM, yielding an EM density map with an overall resolution of 4.5 Å (see Methods and Extended Data Figs. 1-4). Unambiguous density was observed for each of the 5 polypeptides (Fig. 1d). To build an atomic model, we fitted our previously reported X-ray structures into the EM density (Extended Data Fig. 5); for regions of the map where *S. cerevisiae* structures were unavailable, we used structures predicted by AlphaFold2-Multimer (AF)^14,23,40^ (Fig. 1e). In general, both the X-ray and AF structures required minimal adjustment to fit well into the EM density (Extended Data Fig. 5). An exception was the non-CATCHR subunit Sec39, an extended α-solenoid that we modeled into the EM density by rigid body fitting groups of helices. We observed relatively weak EM density for the interacting N-terminal regions of the two CATCHR subunits Dsl1 and Tip20, but this density was nevertheless consistent with the AF prediction (Extended Data Fig. 5d). Finally, no interpretable EM density was observed for three segments of the complex: the 111-residue Dsl1 loop known as the lasso and the C-terminal regions, including the SNARE motifs, of both SNAREs. Importantly, as discussed below, these three segments mediate membrane tethering.

The Dsl1:Qb:Qc model is a pyramidal tower about 25 nm tall (Fig. 2). The broad base of the tower is formed by CATCHR domains B-E of Tip20, the NTDs of the two SNAREs, and the N-terminus of Sec39, with Tip20 and the NTDs oriented roughly perpendicular to the long axis of the overall complex. The α-solenoid subunit Sec39 rises from this base nearly to the top of the complex, where it forms a T-junction with the C-terminal half (CATCHR domains C-E) of the Dsl1 subunit^23^. Notably, this portion of the Dsl1 subunit contains both of the elements that have been implicated in vesicle capture: the lasso (an insertion into domain C) and domain E^28,41–43^. Thus, the Sec39 subunit connects the SNARE-binding end of the Dsl1:Qb:Qc complex to the COPI vesicle-binding end. Its functional importance is underscored by the previous observation that mutations that disrupt the Sec39:Dsl1 T-junction are lethal in yeast^23^.

**Fig. 2:**
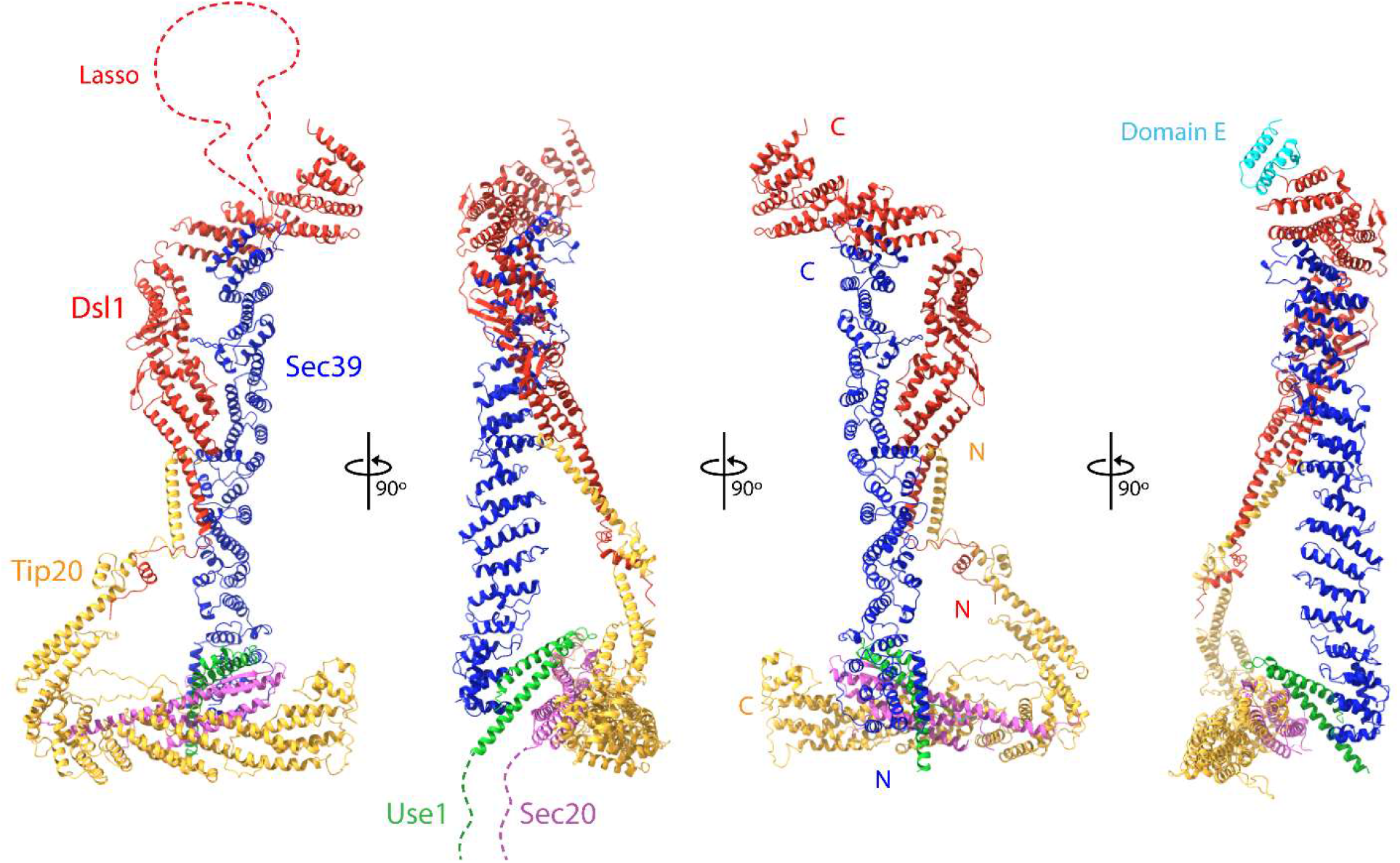
Structure of Dsl1:Qb:Qc. Multiple views of the Dsl1:Qb:Qc complex. Dashed lines represent the lasso (Dsl1 residues 378-488) and the C-termini of two SNARE proteins Sec20 (residues 184-275) and Use1 (residues 86-212).

The two ends of the Dsl1:Qb:Qc complex are also bridged by the interaction between the N-terminal regions of Tip20 and Dsl1 (Fig. 2). The Tip20:Dsl1 bridge is not essential, as the entire N-terminal region of Tip20 can be deleted without compromising yeast growth^14,23^. It becomes essential, however, when the Sec39:Dsl1 interaction is compromised; mutations that weaken the Tip20:Dsl1 interaction are synthetically lethal with mutations that weaken the Sec39:Dsl1 T-junction^23^. Weaker EM density, as well as 3D Variability Analysis^44^, both indicate that the Tip20:Dsl1 bridge is intrinsically flexible (Supplementary Video 1). This flexibility is, however, greatly constrained by the remainder of the Dsl1:Qb:Qc complex, explaining why Dsl1:Qb:Qc is far less conformationally heterogeneous than the Dsl1 complex alone.

### A non-canonical SNARE-SNARE interaction

Among the best-resolved elements in the Dsl1:Qb:Qc EM density are the Qb- and Qc-SNARE NTDs (Fig. 3a). The NTD of the Qb-SNARE Sec20 adopts the 3-helical Habc fold found in the NTDs of many Qa-, Qb-, and Qc-SNAREs including *Eremothecium gossypii* Sec2 0^38,39,45–49^ (Fig. 3b). Compared to these structures, *S. cerevisiae* Sec20 has a novel feature: a pair of antiparallel β-strands between Hb and Hc. The NTD of the Qc-SNARE Use1 forms a modified Habc domain that lacks an N-terminal Ha helix (Fig. 3b). This ‘Hbć fold has not previously been observed; indeed, even the orthologous *Kluyveromyces lactis* Use1 has a typical Habc domain^39^ (see Methods for a detailed comparison). In any case, the absence of an Ha helix would not appear to influence the subunit interactions we observe in Dsl1:Qb:Qc.

**Fig. 3:**
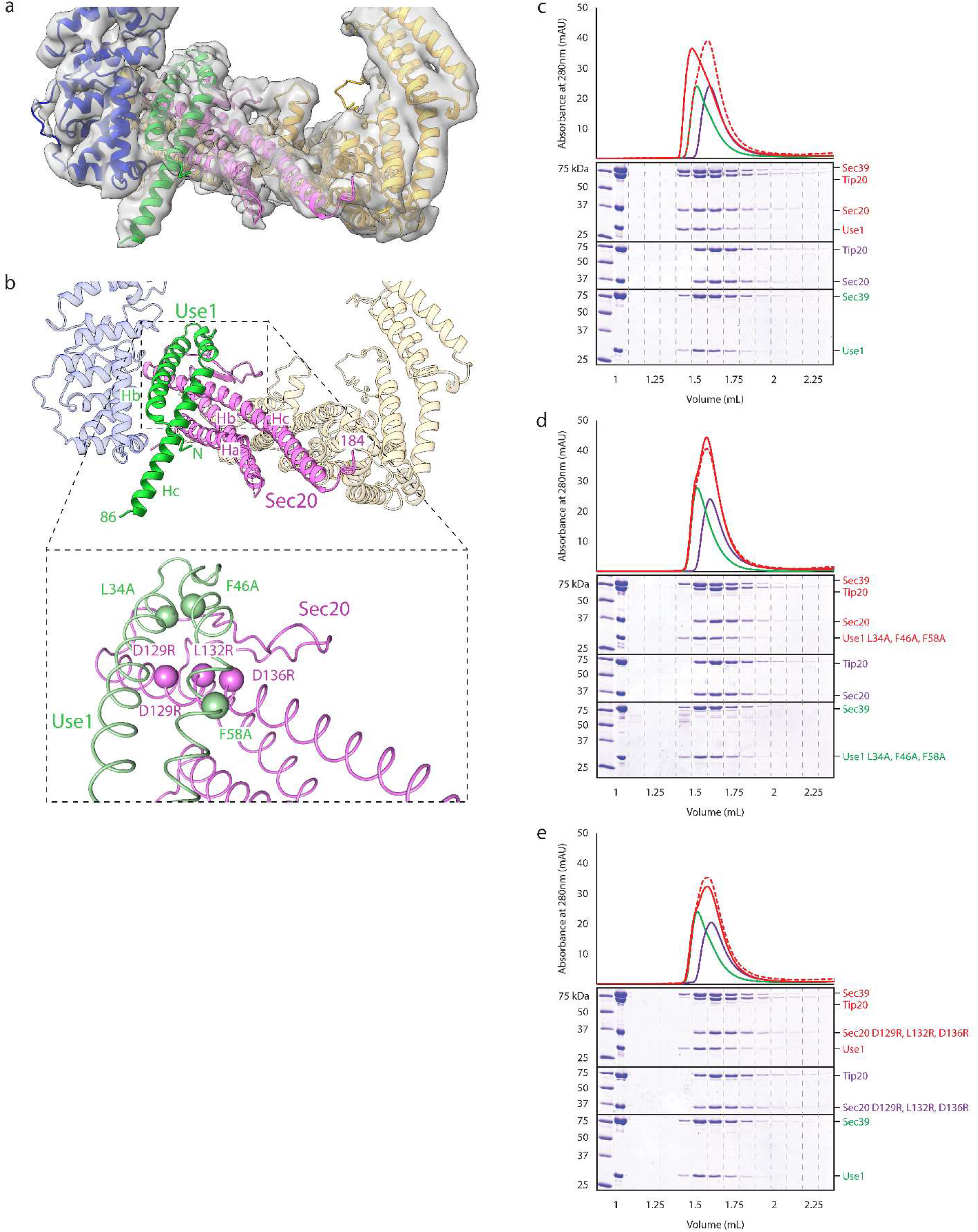
Use1:Sec39 binds Sec20:Tip20 via inter-SNARE interactions. **a,** The broad base of the Dsl1: Qb:Qc complex. Both model and EM density are shown. **b,** SNARE NTDs and their interaction. The Sec20 NTD displays a 3-helix Habc fold, whereas the Use1 NTD displays a non-canonical Hbc fold. Sec39 and Tip20 do not contribute to the SNARE:SNARE interface. The inset shows the location of mutations designed to disrupt the inter-SNARE interaction. **c**, Size-exclusion chromatography (Superdex 200 Increase 3.2/300) was used to analyze Sec20:Tip20 (purple), Use1:Sec39 (green), and a mixture of the two (red). All polypeptides were present at 10 μM final concentration. The dashed red line represents the arithmetic sum of the Sec20:Tip20 and Use1:Sec9 profiles, as expected for non-interacting samples. The large size and elongated shape of both Sec20:Tip20 and Use1:Sec39 may explain the lack of a larger shift in elution volume upon formation of the quaternary complex. **d** and **e**, Experiments were performed as in (c), but with complexes containing either Use1 (L34A, F46A, F58A) or Sec20 (D129R, L132R, D136R). No binding is detected.

Paradigmatically, SNAREs interact via their SNARE motifs^50^. By contrast, the Dsl1:Qb:Qc structure reveals two SNAREs interacting via their NTDs. This NTD:NTD interaction between Sec20 and Use1 links together Tip20 and Sec39, two Dsl1 complex subunits that were previously proposed to function as independently mobile legs^23,39^ (Fig. 3a,b). The NTD:NTD interface, like the other protein:protein interfaces in Dsl1:Qb:Qc, is not well conserved at the sequence level, suggesting rapid co-evolution. Nonetheless, this interaction was recently predicted by a high-throughput computational search for interacting yeast proteins^51^, and its formation in Dsl1:Qb:Qc buries an interfacial surface accessible area of over 1,000 Å^2^.

To validate the Qb-SNARE:Qc-SNARE interaction *in vitro*, we mixed Sec20:Tip20 and Use1:Sec39 and then tested for complex formation using size exclusion chromatography. We used complexes of SNAREs and tethering subunits, rather than Sec20 and Use1 SNAREs by themselves, to ensure that the NTDs were properly folded. Wild-type Sec20:Tip20 bound to wild-type Use1:Sec39, as judged by a noticeable shift to an earlier elution volume compared to Sec20:Tip20 or Use1:Sec39 alone (Fig. 3c). To confirm that binding involves a direct NTD:NTD interaction, we designed triple mutations to disrupt the interface: Sec20 (D129R, L132R, D136R) and Use1 (L34A, F46A, F58A) (Fig. 3b). The mutant SNAREs formed stable complexes with Tip20 or Sec39 (Extended Data Fig. 7) but, as predicted, abolished binding of Sec20:Tip20 to Use1:Sec39 (Fig. 3d,e).

### Cooperative function of the Dsl1:Qb:Qc complex

The Dsl1:Qb:Qc structure implies that the NTDs of the ER SNARE proteins Sec20 and Use1 are not simply a means of anchoring the Dsl1 MTC to the membrane, but are integral structural components of the MTC itself. To investigate the functions of the NTDs *in vivo*, we used plasmid shuffling to replace the wild-type yeast SNAREs with several mutants. We previously reported that deleting the NTD of Sec20 was lethal^39^; correspondingly, we found that deleting the NTD of Use1 was lethal (Fig. 4a). Thus, not only the SNAREs, but their NTDs specifically, are essential for yeast viability, presumably because they are essential for Golgi-to-ER trafficking.

**Fig. 4:**
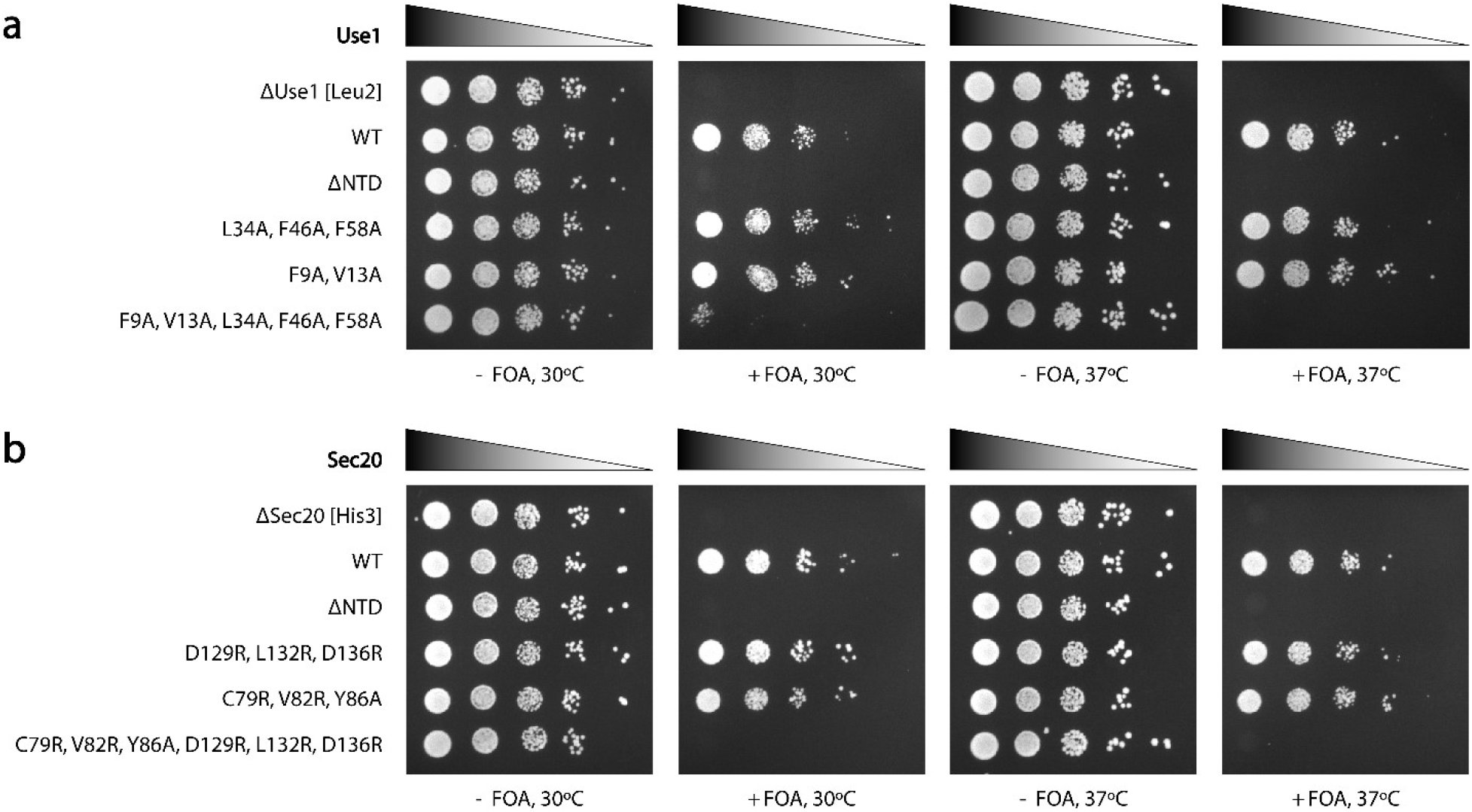
SNARE incorporation into Dsl1:Qb:Qc is essential in yeast. **a**, Yeast strains lacking endogenous Use1 were maintained using a wild-type Use1 covering plasmid marked with Ura3 and a second plasmid with the Leu2-linked Use1 allele indicated at the left. The viability of these alleles is indicated by growth on 5-fluoroorotic acid (FOA) selective plates, which leads to the loss of the covering plasmid. **b**, Yeast strains lacking endogenous Sec20 were maintained using a wild-type Sec20 covering plasmid marked with Ura3 and a second plasmid with the His3-linked Sec20 allele indicated at the left. The viability of these alleles is indicated by growth on 5-fluoroorotic acid (FOA) selective plates, which leads to the loss of the covering plasmid.

To probe the importance of the NTD:NTD interface, we replaced each SNARE in turn with the triple mutants described above, Sec20 (D129R, L132R, D136R) and Use1 (L34A, F46A, F58A). Surprisingly, neither replacement caused a major growth defect (Fig. 4). Next, we designed mutations to disrupt the Sec20:Tip20 and Use1: Sec39 interfaces: Sec20 (C79R, V82R, Y86A) and Use1 (F9A, V13A). Consistent with our design goal, mutant Sec20 failed to co-purify with bacterially co-expressed Tip20, and mutant Use1 failed to co-purify with bacterially coexpressed Sec39 (Extended Data Fig. 8). Each mutant protein was introduced via plasmid shuffling into yeast and again was able to support apparently wild-type growth (Fig. 4). Although this, too, was surprising, it was consistent with our previous finding that yeast tolerated a mutation in Tip20, (I481D, L585D), that lowers its affinity for Sec20 at least 15-fold^39^. Thus, it was possible to compromise the Sec20:Use1, Sec20:Tip20, or Use1:Sec39 interfaces without markedly affecting yeast growth; these findings are further discussed below. Finally, we combined the Sec20 mutations to generate a Qb-SNARE unable to bind either Use1 or Tip20. Notably, this combination was lethal. Also lethal was the combination of Use1 mutations to generate a Qc-SNARE unable to bind either Sec20 or Sec39 (Fig. 4). Thus, mutant Qb- and Qc-SNAREs that cannot incorporate into the Dsl1:Qb:Qc complex, either because they lack an NTD or because they bear NTD mutations that disrupt both interfaces with the remainder of the complex, cannot support yeast viability. Conversely, it appears that each SNARE can be functionally incorporated into the Dsl1:Qb:Qc complex in two different ways, either by binding to its partner Dsl1 complex subunit or by binding to the other SNARE.

### Dsl1: Qb:Qc resembles exocyst

A side-by-side comparison of the Dsl1:Qb:Qc and exocyst complexes reveals striking similarities, despite their very different subunit compositions and internal architectures (Fig. 5). Each complex is a roughly pyramidal tower 25-30 nm in height, although the exocyst tower is broader owing to its much greater molecular weight^11^. The wide base of each complex is defined by a single CATCHR-family subunit – Tip20 for the Dsl1 complex, Sec6 for exocyst – oriented approximately perpendicular to the long axis of the complex. In each complex, this base subunit binds directly to a SNARE present on the target membrane: Tip20 binds to the ER Qb-SNARE Sec20, which in turn binds to the Qc-SNARE Use1, whereas in exocyst Sec6 binds to the plasma membrane Qb/Qc SNARE Sec9^52,53^. Finally, the distal tip of each complex is implicated in vesicle capture. As noted above, the Dsl1 complex binds COPI vesicles using the lasso, and probably the E domain, of the Dsl1 subunit^41,42,54^. Exocyst binds the secretory vesicle Rab protein Sec4 using its Sec15 subunit^55^; studies of *Drosophila melanogaster* Sec15 suggest that CATCHR domain D, situated at the tip of the complex, contains the Rab binding site^11,16^. Thus, key aspects of the two complexes, including their shapes and overall dimensions, as well as the relative dispositions of important vesicle and target membrane binding sites, are remarkably congruent.

## DISCUSSION

The Dsl1 complex, with only 3 subunits, is a minimalist MTC. Its SNARE-binding subunits Tip20 and Sec39 had been presumed, based on negative-stain EM, to function as independent legs, with this flexibility potentially important for its tethering function^23^. Similarly, negative-stain EM studies of two additional CATCHR-family complexes, GARP and COG, imply the presence of multiple, flexible legs^56,57^. Against this backdrop, two aspects of our findings are especially notable. First, when the Dsl1 complex is bound to Qb- and Qc-SNAREs – as it would be on the ER membrane – it adopts a closed conformation mediated by a novel, direct interaction between the SNARE NTDs (Fig. 2). Second, this SNARE-dependent – and thus membrane anchoring-dependent – closed conformation bears a strong resemblance to exocyst, the only other CATCHR-family complex with a known structure^11^ (Fig. 5). Both Dsl1:Qb:Qc and exocyst are 25-30 nm long, with binding sites for target membrane SNAREs at one end and for vesicle proteins at the other end. Another common feature is a hole of similar dimensions near the broad base of each complex (Fig. 5). Intriguingly, this hole is greatly enlarged in a gain-of-function exocyst mutant (Exo70 I114F)^58^; its functional significance, however, is unclear. Overall, the convergent properties of the two very different CATCHR-family complexes suggest that these properties are essential for membrane tethering and fusion.

The Dsl1 complex contains a single pair of CATCHR-family subunits, Tip20 and Dsl1, which interact in the canonical antiparallel manner via their N-terminal ends^14^. Their CATCHR domains, meanwhile, bind Sec20 and COPI, respectively^39,42^. Thus, a single pair of interacting CATCHR-family subunits satisfies the minimal requirement for any tethering factor: binding two membranes simultaneously (Fig. 6). The non-CATCHR Sec39 subunit is nonetheless critical, helping recruit SNAREs and thereby generating a stable complex with a well-defined conformation. Exocyst, by contrast, has a far more elaborate architecture, with four different pairs of interacting CATCHR-family subunits^11,59^. None of these pairs, however, links one membrane to the other. Indeed, the SNARE-binding subunits, Sec3 and Sec6, reside within one 4-subunit subcomplex, whereas the Rab-binding Sec15 subunit resides within the other^20^. Thus, the tethering activity of exocyst appears to depend on the structural integrity of the entire complex.

**Fig. 5:**
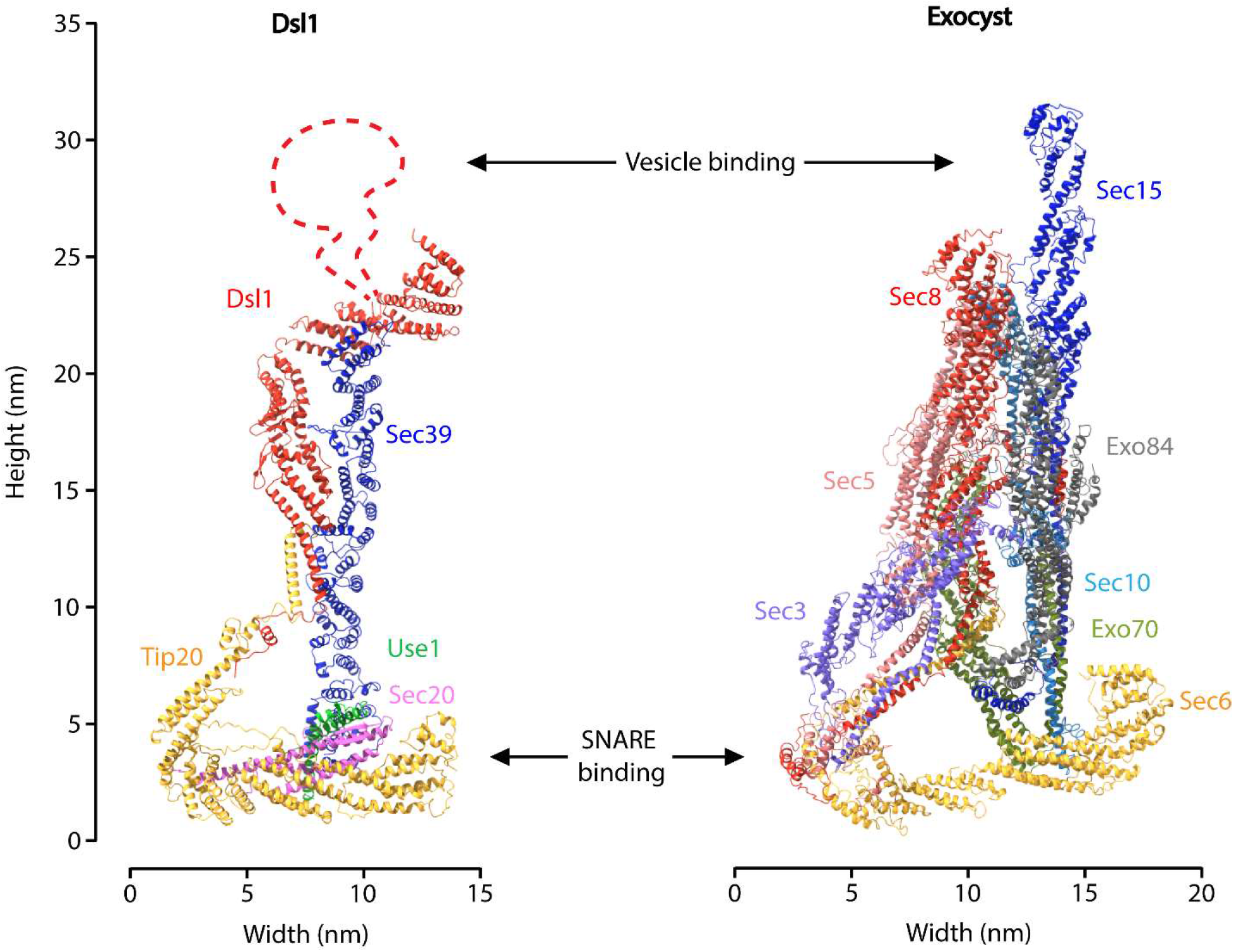
Comparison of Dsl1:Qb:Qc and exocyst. Cryo-EM structures of Dsl1:Qb:Qc and exocyst (PDB code 5YFP) are compared side by side. The N-terminal half of exocyst subunit Sec3, which contains a PH domain flanked by long sequences that are predicted to be disordered, could not be modeled into the cryo-EM map^11^. Chemical crosslinking suggests that this PH domain, which binds the Qa-SNARE Sso1/2, is located near the Sec6 subunit^70,72^. Sec6 bind the R-SNARE Snc2, and SNARE complexes, in addition to the Qb/Qc-SNARE Sec9^52,73^.

The straightforward architecture of the Dsl1:Qb:Qc complex, in which each subunit interacts with two neighbors in a cyclical arrangement (Fig. 2), appears to confer functional robustness. Whereas all 5 subunits are essential, 4 of the 5 inter-subunit interfaces (the exception being Dsl1:Sec39) can be destabilized, one at a time, without compromising yeast growth (Fig. 4 and refs. ^14,23,39^). This functional robustness presents a conundrum. On the one hand, the stability of the closed conformation, and its striking resemblance to exocyst, strongly imply that it is functionally relevant. On the other hand, interface mutants likely to destabilize this conformation are tolerated in vivo. One potential resolution to this conundrum is that the closed conformation, while functionally important, is not rate-limiting for yeast growth under standard laboratory conditions. Another possibility is that other trafficking factors, such as the cognate Qa-SNARE Ufe1, R-SNARE Sec22, and/or Sec1/Munc18- (SM-) family protein Sly1, are capable of stabilizing the closed conformation of mutation-bearing complexes at functionally critical junctures. Structures incorporating these factors may be needed to address this issue more fully.

Our results show that integration of both SNAREs into the Dsl1:Qb:Qc complex is essential for yeast viability, since mutations that expel either SNARE by deleting or mutating its NTD are lethal. It therefore seems likely that the Dsl1:Qb:Qc complex plays an important role in scaffolding SNARE assembly, as previously proposed^14,23,39^. The two SNARE motifs are not visible in the EM density, consistent with the expectation that they are natively unfolded in the absence of the Qa- and R-SNAREs. Nevertheless, their co-incorporation into a Dsl1:Qb:Qc complex brings the two SNARE motifs into relative proximity, by virtue of the 36- and 74-residue linkers that connect them to their NTDs. This general proximity stands in contrast with the precise alignment of Qa- and R-SNAREs by SM proteins, which function as templates for SNARE assembly^60,61^. It will be important in future work to elucidate how the SNAREs in Dsl1:Qb:Qc assemble with their cognate Qa- and R-SNAREs – presumably with the assistance of the SM protein Sly1 – to generate a fusogenic complex.

Although CATCHR and HOPS/CORVET-family MTCs were long thought to be conformationally variable, single-particle cryo-EM studies of exocyst^11^, HOPS^62^, and now Dsl1:Qb:Qc have revealed relatively rigid cores with widely separated membrane-binding sites. It may instead be the attachments formed between MTCs and membranes that are flexible. This flexibility is exemplified by the unfolded SNARE regions that anchor the Dsl1:Qb:Qc complex to the ER and the disordered lasso that captures COPI vesicles (Fig. 6). Many other MTCs, including exocyst and HOPS, engage membranes by binding to Rab proteins. This mode of attachment is also likely to be flexible, since Rabs are anchored to membranes by ~30-40-residue C-terminal hypervariable regions^63^. Thus, despite drastic differences in internal architecture, 3 MTCs from 2 different families reveal relatively rigid scaffolds with widely separated sites for flexible membrane attachments. While these properties cannot themselves explain how membranes are brought together, they are likely to be essential for tethering, membrane curvature formation, and subsequent fusion.

**Fig. 6:**
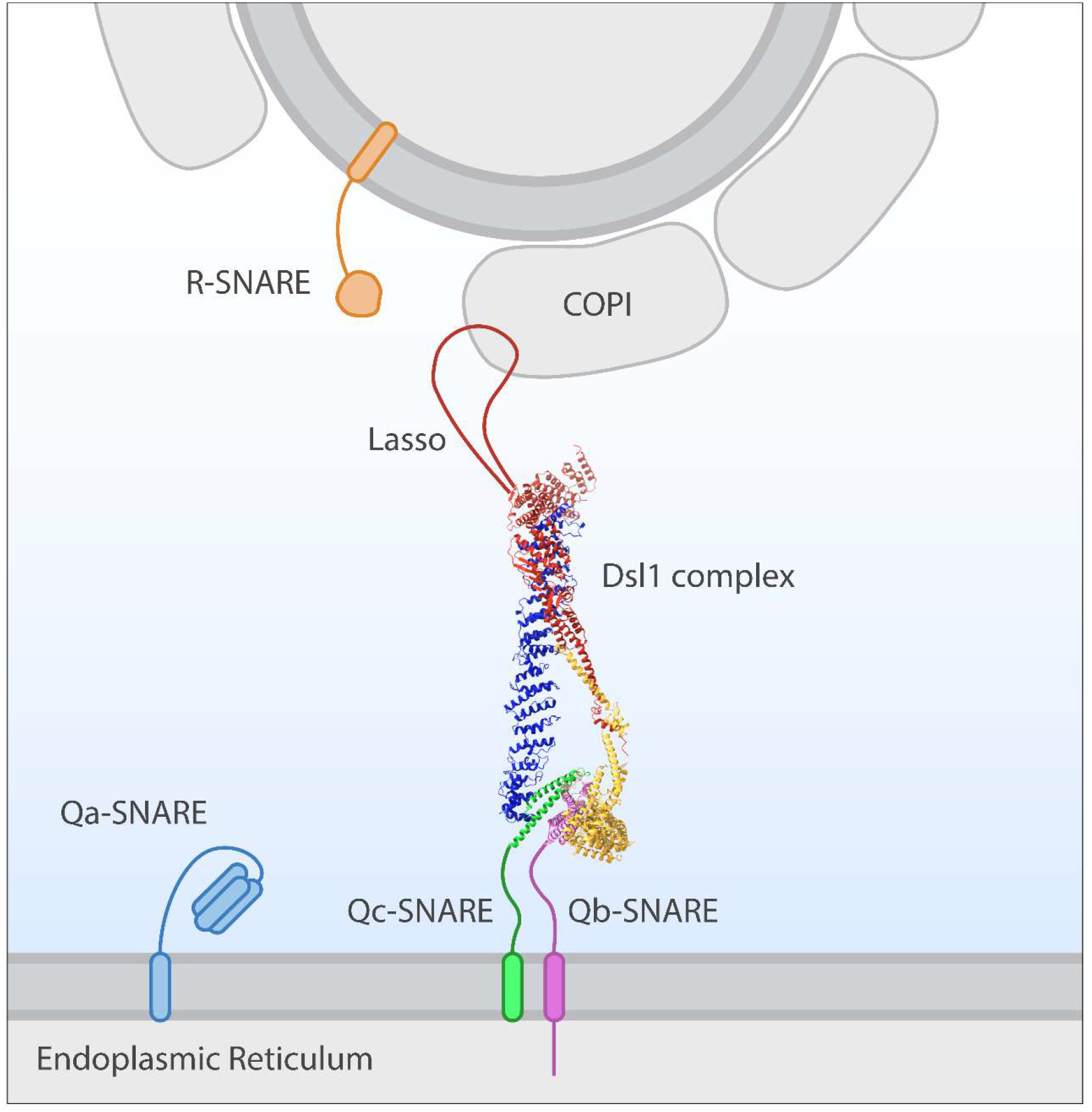
Model for retrograde vesicle capture by Dsl1: Qb:Qc.

## ACKNOWLEDGMENTS

We thank Xiao Fan, Paul Shao, Venu Vandavasi, and members of the Hughson laboratory past and present for helpful advice and discussion. We are grateful to the Princeton University Biophysics and Macromolecular Crystallography core facilities for technical assistance. We acknowledge the use of Princeton’s Imaging and Analysis Center (IAC), which is partially supported by the Princeton Center for Complex Materials (PCCM), a National Science Foundation (NSF) Materials Research Science and Engineering Center (MRSEC, DMR-2011750). This work was supported by National Institutes of Health grants R01GM071574 (FMH), T32GM007388 (KAD, AES, JDS, SMT), and F31GM12676 (SMT).

## METHODS

### Protein Expression and Purification

The complete *S. cerevisiae* Dsl1:Qb:Qc complex, as well as Tip20:Qb and Sec39:Qc subcomplexes, were overproduced using pQLink-based co-expression plasmids^64^. All co-expression plasmids were generated by concatenating pQLink plasmids that each bore a single open reading frame derived from yeast genomic DNA. These open reading frames corresponded to the Dsl1 complex subunits Dsl1, Tip20, or Sec39, the cytoplasmic region (residues 1-275) of the Qb-SNARE Sec20, or the cytoplasmic region (residues 1-212) of the Qc-SNARE Use1. N-terminal or C-terminal heptahistidine (His_7_) tags were added using pQLinkH (Addgene plasmid 13667) or a modified derivative; otherwise, we used pQLinkN (Addgene plasmid 13670). The Dsl1:Qb:Qc complex was overproduced using a single concatenated pQLink plasmid bearing genes encoding His_7_-Dsl1, Sec39, Tip20, Sec20, and Use1. For biochemical studies, we generated two plasmids, one for overproducing Tip20:Sec20-His_7_, the other for overproducing Sec39:His_7_-Use1. For pull-down experiments, we generated plasmids for overproducing His_7_-Tip20:Sec20 and His_7_-Sec39:Use1. Mutations were introduced using the Q5 Site-Directed Mutagenesis Kit (NEB). All plasmids were validated by DNA sequencing.

Protein complexes were overproduced in BL21-CodonPlus (DE3) *Escherichia coli* cells (Agilent) grown in high salt LB Broth (Sigma) at 37°C to an OD_600_ of 0.5-0.7 and induced by the addition of 0.5 mM isopropyl β-D-1-thiogalactopyranoside. The cells were harvested after 16 h of induction at 15°C, resuspended in 300 mM NaCl, 20 mM HEPES pH 7.5 (HBS), supplemented with 5 mM β-mercaptoethanol, 1 mM phenylmethylsulfonyl fluoride, and 10 mg/ml DNase I (Roche), and lysed using a cell disrupter (Avestin). After clarifying the lysate via centrifugation, protein complexes were purified from by Ni^2+^-NTA (Takara Bio) affinity chromatography (Dsl1:Qb:Qc and His_7_-Tip20:Sec20) or TALON (Takara Bio) affinity chromatography (all other complexes).

Proteins for electron microscopy or *in vitro* binding assays were further purified by anion exchange and size-exclusion chromatography (MonoQ and Superdex 200 Increase, Cytiva). Purified proteins were concentrated, flash frozen, and stored at −80°C in 150 mM NaCl, 20 mM HEPES pH 7.5 supplemented with either 1 mM (for electron microscopy) or 5 mM (for *in vitro* binding assays) dithiothreitol.

### Electron Microscopy and Data Processing

For negative stain microscopy, protein was diluted to a concentration of 25 nM and applied to CF400-copper ultra-thin support carbon grids (Electron Microscopy Sciences) that were glow discharged with a PELCO easiGlow (Ted Pella, Inc). Staining was performed by applying 0.75% (w/v) uranyl formate and immediately removing by blotting with filter paper. Application was repeated five times, with a 30 s room temperature incubation before the final blotting step. Images were collected at a pixel size of 2.02 Å with a Talos F200X Scanning/Transmission Electron Microscope (Thermo Fisher Scientific) at 200 kV. 6,013 particles were manually picked and sorted into 25 2D-class averages utilizing RELION v3.0.6^65^. For cryogenic electron microscopy, protein was diluted to a concentration of 10 μM in storage buffer supplemented with 0.05% (v/v) NP40, applied to Quantifoil R 1.2/1.3 Cu300 grids (Electron Microscopy Sciences) glow discharged using a NanoClean Model 1070 (Fischione) for 30 s, and frozen in liquid ethane with a Vitrobot Mark IV (Thermo Fisher Scientific) at 100% humidity and 4°C with a blot time of 6 s. Particles were imaged in a Titan Krios G3i Cryo Transmission Electron Microscope (Thermo Fisher Scientific) at 300 kV with a K2 Summit direct electron detector camera with GIF Quantum energy filter (Gatan). Images were collected with the software EPU in EFTEM mode with a pixel size of 1.114Å, defocus range of −1.25 to −2.5 μm, exposure time of 5.6 s, and total electron dose per image of 45 e^-^/Å^2^ (Extended Data Fig. 1). Motion correction and all subsequent processing steps were performed using cryoSPARC v.4.1.1^66^. 469,193 particles were template-picked from 5,857 collected micrographs utilizing a low-resolution density map of the complex generated from a subset of initial micrographs. Next, 50 2D class averages were generated and 26 classes, containing 286,801 particles, were selected to proceed with. The total number of particles was further reduced by generating multiple sets of 3D class averages and proceeding using the most promising class (or a combination of multiple classes from the same set, if that resulted in the highest-resolution map at that step). This step was repeated three times, reducing the total number of particles to 49,947. Initial refinement using the Non-Uniform Refinement tool in cryoSPARC produced a 6.2 Å-resolution map (Extended Data Fig. 2). Additional local refinement was performed on four overlapping ~150 kDa segments of the complex, resulting in a 4.5 Å-resolution composite map (Extended Data Fig. 3). Local resolution was generated with the Local Resolution Estimation tool in cryoSPARC and resolution was presented using GSFCS scores (Extended Data Fig. 4). Local refinement outputs were combined in ChimeraX^67^ for model building and figure generation. An intermediate, 7.2 Å-resolution map was utilized for 3D Variability Analysis (3DVA)^44^, also in cryoSPARC (Supplementary Video 1). This map was chosen due to the relative lack of noise in the 3DVA output.

### Model Building

To construct an atomic model of the Dsl1:Qb:Qc complex, we used published *S. cerevisiae* crystal structures whenever they were available. Tip20 (PDB code 3FHN), as well as Dsl1 residues 37-355 (3ETU), were rigid body fitted into the EM density domain by domain with only minimal adjustments needed. Sec39 (3K8P) lacks the domain architecture of Dsl1 and Tip20, so it was instead modeled into the EM density by rigid body fitting groups of helices. Together, these crystal structures covered more than 70% of the final Dsl1:Qb:Qc model (Extended Data Fig. 5a).

To build the remainder of the model, we used structures predicted using AlphaFold2^68^ and AlphaFold-Multimer^40^ (Extended Data Figs. 5a and 6). The predicted structure of Dsl1 residues 356-754 fit the EM density well (correlation coefficient of 0.901; Extended Data Figs. 5b and S6a-b) except for the “lasso”, residues 378-488. The lasso was omitted from the model based on its low sequence complexity, predicted disorder based on high per-residue pLDDT scores, and lack of corresponding EM density. Sec39 residues 1-112 was generated using Alphafold2, since the corresponding region of the published *S. cerevisiae* crystal structure (3K8P) displayed weak electron density and lacked residues 1-29 and 63-100 altogether (correlation coefficient of 0.891; Extended Data Figs. 5c and 6a,c). The X-ray structure also lacks several loops which were supplied using the structure predicted by AlphaFold2. Finally, we used AlphaFold-Multimer to generate a model of the Dsl1:Tip20 interface (Dsl1 residues 1-131 and Tip20 residues 1-66). The resulting model predicts that Dsl1 residues 1-36 bind to Tip20. Although the local resolution of the EM density is relatively low (approximately 9 Å), the model generated by AlphaFold-Multimer fits reasonably well (correlation coefficient of 0.736; Extended Data Figs. 5a and 6a,d).

We used AlphaFold-Multimer to model the SNAREs, Sec20 and Use1. The predicted structure of the complex of the two NTDs (Sec20 residues 1-184 and Use1 residues 1-86) fit remarkably well into the EM density map, with no significant regions of disagreement (correlation coefficient of 0.858; Extended Data Figs. 5a and 6a,e). Fitting the SNAREs individually into the EM density yielded almost indistinguishable results. No EM density was observed for more C-terminal regions of the SNAREs or for Sec20 residues 38-51.

Following initial building, the fit of the model to the EM map was optimized using phenix.real_space_refine (version 1.17)^69^ with restraints on secondary structure and rotamers. The CC(mask) between the model and the EM map was 0.783.

### Comparisons with X-ray structures

We previously reported the 6.5 Å-resolution X-ray structure of *Kluyveromyces lactis* Use1 bound to Sec39 and a 306-residue C-terminal fragment of Dsl1^39^ (PDB code 6WC4). In that structure, *K. lactis* Use1 was tentatively modeled in the reverse N-to-C orientation compared to *S. cerevisiae* Use1 in the Dsl1: Qb:Qc structure reported here. To investigate this discrepancy, we used AlphaFold-Multimer to predict the heterodimeric complex of *K. lactis* Use1 and Sec39. For rigid body fitting to the original electron density map, the predicted structure was split in two at Sec39 residue 366, yielding a model that agreed well with the map. In this model, *K. lactis* Use1 adopts the same N-to-C orientation as *S. cerevisiae* Use1, and its Hb and Hc helices occupy the same positions (relative to Sec39) as the Hb and Hc helices of *S. cerevisiae* Use1 (Extended Data Fig. 5e). We conclude that *S. cerevisiae* Use1 lacks the Ha helix but otherwise has the same overall structure as *K. lactis* Use1. Moreover, the Sec39:Use1 binding mode is conserved. The PDB entry for the *K. lactis* Use1:Sec39:Dsl1 complex has been updated accordingly (new PDB code 8FTU).

Previously, in an effort to characterize the interface between Tip20 and Dsl1, we obtained the crystal structure of a fusion protein that connected *S. cerevisiae* Tip20 residues 1-40 to Dsl1 residues 37-339 via a short linker^14^. That structure, together with structure-based mutagenesis, provided evidence that the Tip20:Dsl1 interaction was mediated by an antiparallel interaction between α-helices near the N-terminus of each subunit. That interpretation is generally confirmed by the Dsl1:Qb:Qc structure, but there are differences. First, one of the two α-helices is shifted by about two helical turns relative to the other. Second, the Tip20:Dsl1 interface is more extensive than anticipated, involving regions of both Tip20 and Dsl1 that were missing from the fusion protein studied previously. Nonetheless, the mutagenesis results obtained previously^14^ are fully consistent with the new structure and reinforce the conclusion that the helix-helix interaction is essential for the stability of the Tip20:Dsl1 interface.

### *In vitro* Binding Assays

For size-exclusion chromatography, proteins were mixed at a final concentration of 10 μM each in a volume of 50 μL 150 mM NaCl, 20 mM HEPES pH 7.5, 5 mM dithiothreitol. After a 5 h incubation at 4°C, samples were analyzed using an Superdex 200 Increase 3.2/300 column (Cytiva). Fractions were visualized using 10% SDS-PAGE polyacrylamide gels.

### Yeast methods

Haploid *S. cerevisiae* strains bearing deletions of either Sec20 or Use1 were maintained with pRS416-derived covering plasmids containing the missing genes. These plasmids, as well as the others used in these experiments, contained the coding regions of Sec20 or Use1 along with 500 bases of upstream flanking DNA and, in the case of Use1, 500 bases of downstream flanking DNA. For plasmid shuffling experiments involving Sec20, yeast were transformed with pRS413 containing either wild-type or mutant Sec20 and plated on synthetic complete agar lacking histidine (to select for pRS413) and uracil (to select for pRS416). Transformants were grown overnight at 30°C in synthetic complete media lacking histidine and diluted to an OD_600_ of 0.2. To select against the covering plasmid, ten-fold serial dilutions were plated onto synthetic complete agar lacking histidine and containing 0.1% w/v 5-fluoroorotic acid (GoldBio). Plasmid shuffling experiments involving Use1 were conducted in an analogous manner. However, because wild-type or mutant Use1 was carried on pRS415 instead of pRS413, we used media lacking leucine instead of media lacking histidine.

