## Extended Data for "Structure of a Membrane Tethering Complex Incorporating Multiple SNAREs"

**Extended Data Figures**

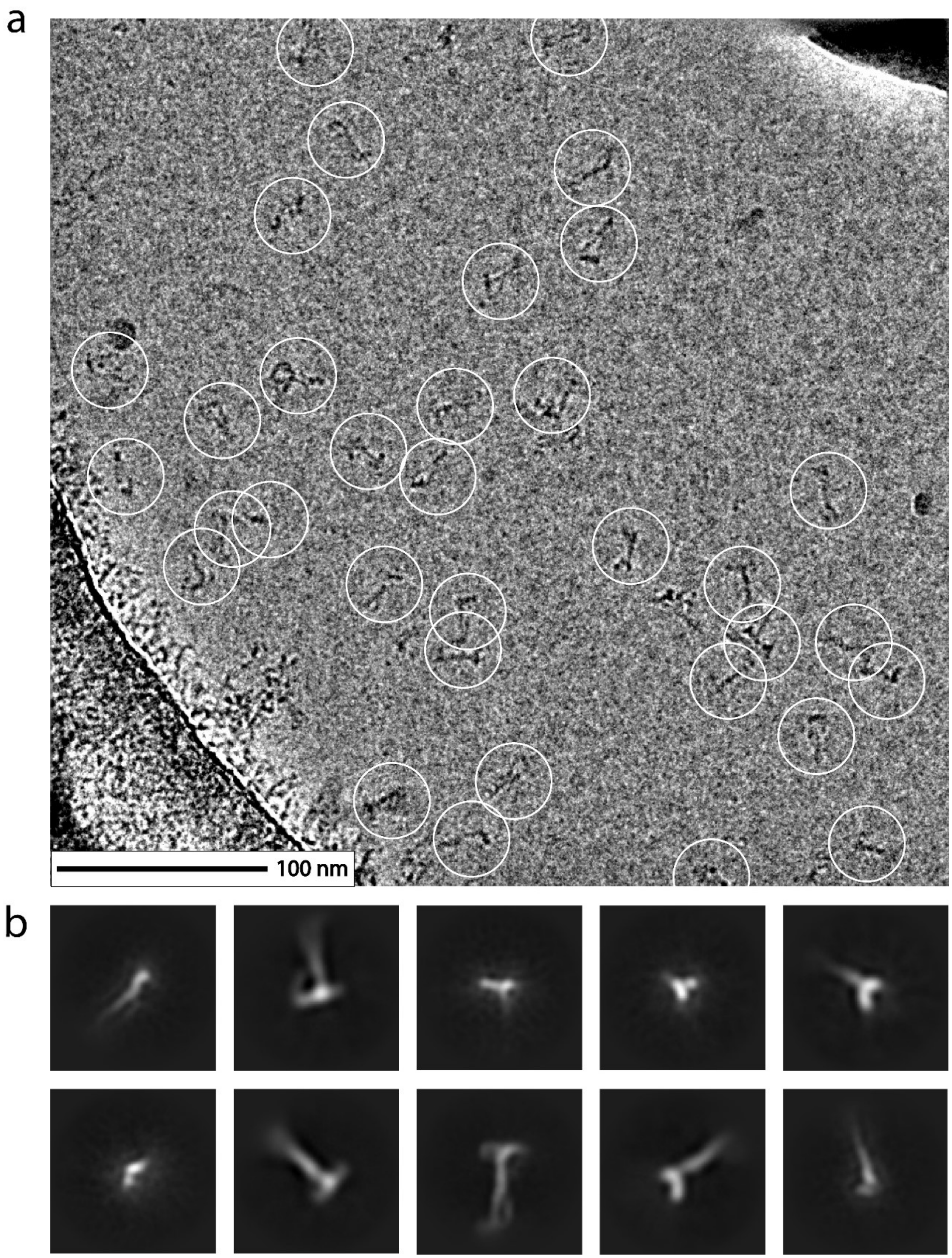


**Extended Data Fig. 1: Representative cryo-EM images. a**, Representative cryo-EM micrograph of Dsl1:Qb:Qc prepared as described in Methods. Individual Dsl1:Qb:Qc particles have been marked with white circles. **b**, Representative 2D class averages of the Dsl1:Qb:Qc complex used in the construction of the cryo-EM density. Classes were generated using cryoSPARC.

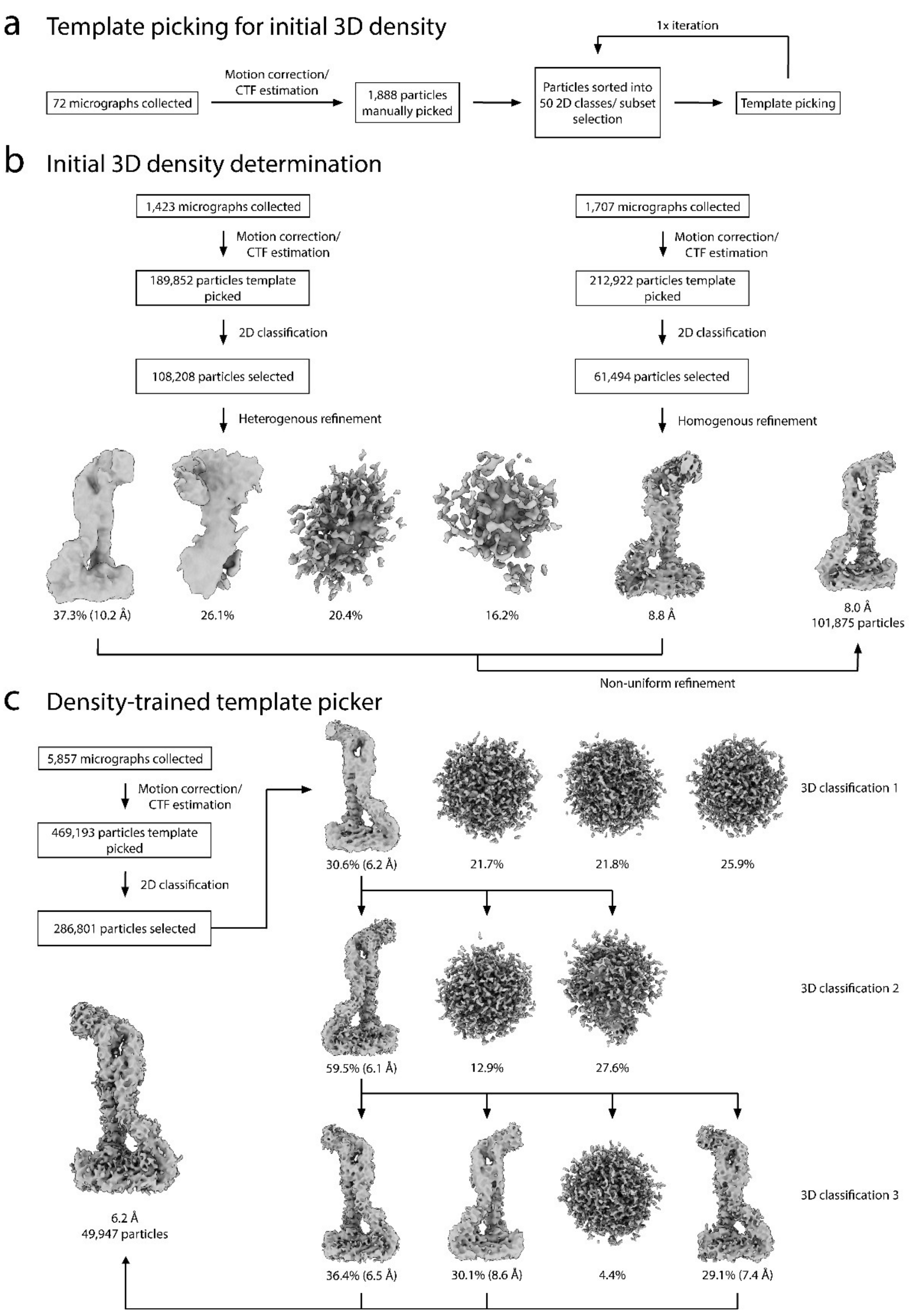


**Extended Data Fig. 2: Workflow of cryo-EM data processing pre-local refinement. a**, Flowchart describing the training of the template picker used in the initial EM map determination of the Dsl1:Qb:Qc complex. **b**, Flowchart describing the method for generating an initial 3D model of Dsl1:Qb:Qc to use for density-guided template picking. **c**, Flowchart describing the method for generating a pre-local-refinement EM map of Dsl1:Qb:Qc using a template picker trained on the 8.0 Å map generated in b. The EM map is inverted relative to the final map.

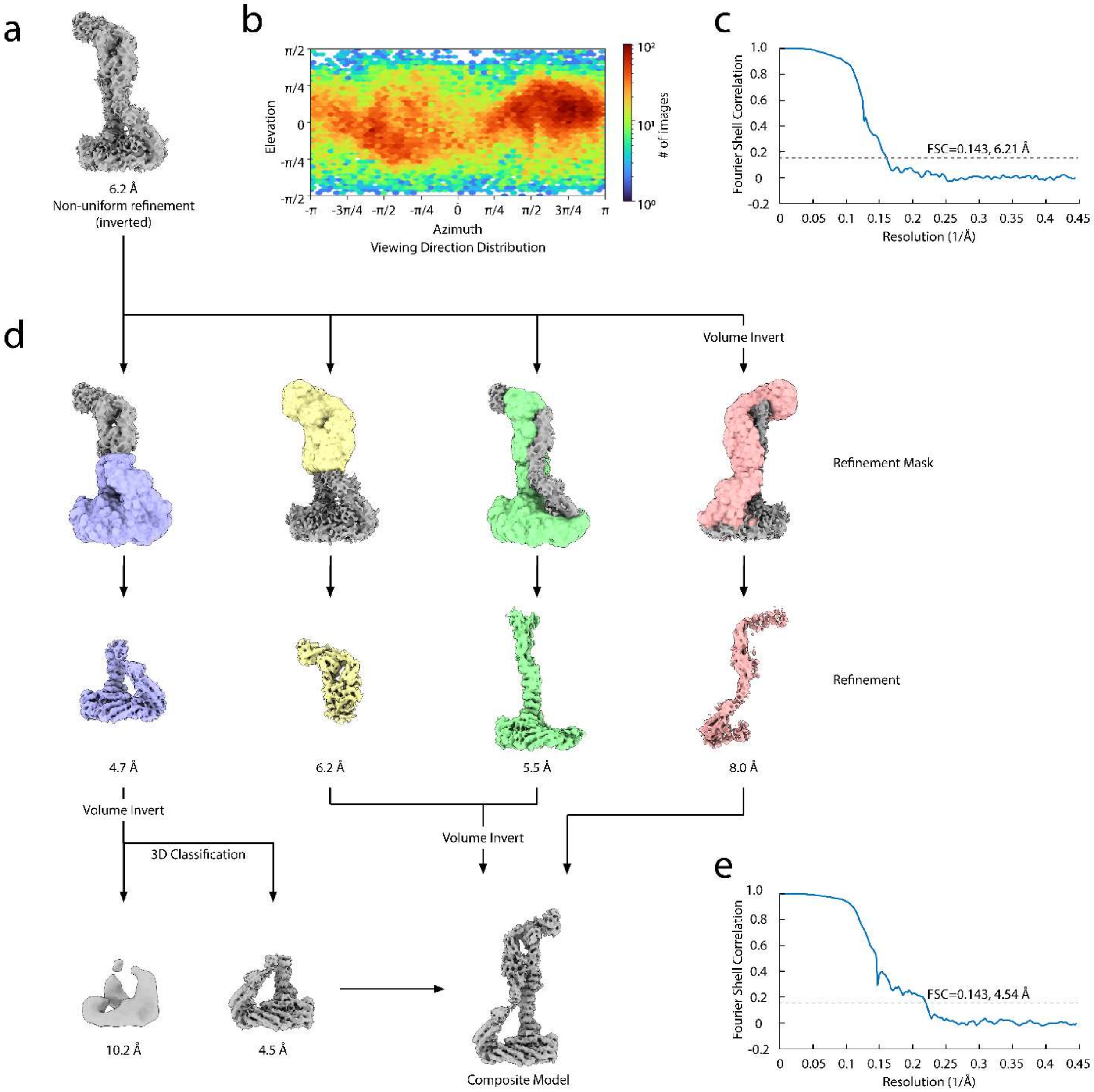


**Extended Data Fig. 3: Local refinement of the Dsl1:Qb:Qc complex. a**, Output from Non-Uniform Refinement that was used for mask generation and local refinement. **b**, Angular distribution of the consensus map. **c**, GS-FSC curve of the consensus map. **d**, Flowchart describing the masking process for local refinement of the EM map. Four separate masks were applied, covering approximately half of the complex in four different orientations. At the outset of this process, the EM map was inverted by non-uniform refinement. Local refinement was performed on both the inverted and corrected EM map, and the higher-resolution output was used in the final composite map. **e**, GS-FSC curve of the composite EM map of the Dsl1:Qb:Qc complex. The curve was generated by combining the half maps from each of the four local refinement jobs and processing with the Validation (FSC) tool in cryoSPARC.

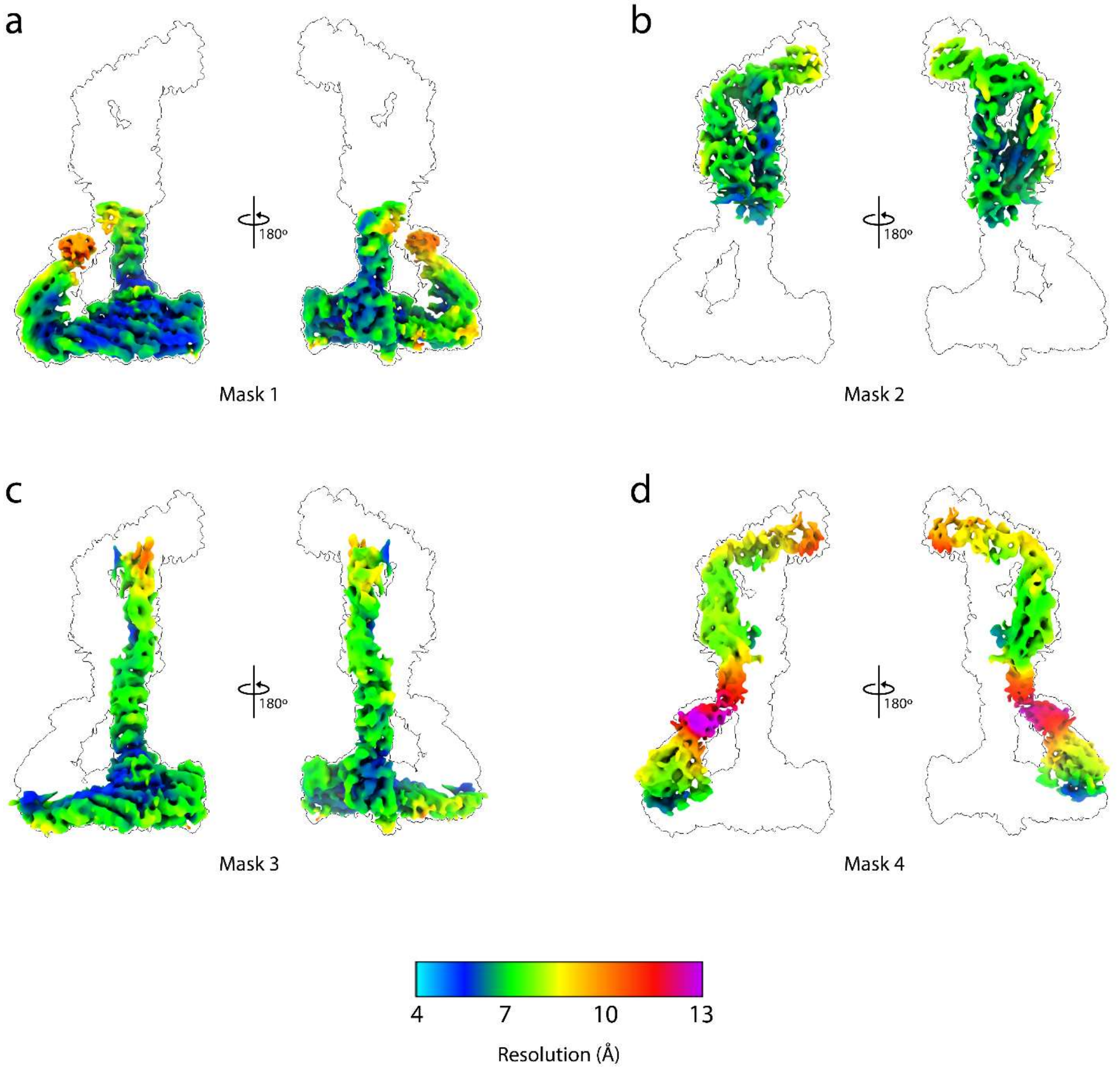


**Extended Data Fig. 4: Local resolution of the Dsl1:Qb:Qc complex cryo-EM map. a**, Local refinement map generated from Mask 1 colored by local resolution from two different viewing angles. The map is superimposed on an outline of the complete Dsl1:Qb:Qc complex at a lower contour level for reference. **b**, Local refinement map generated from Mask 2 colored by local resolution. **c**, Local refinement map generated from Mask 3 colored by local resolution. **d**, Local refinement map generated from Mask 4 colored by local resolution.

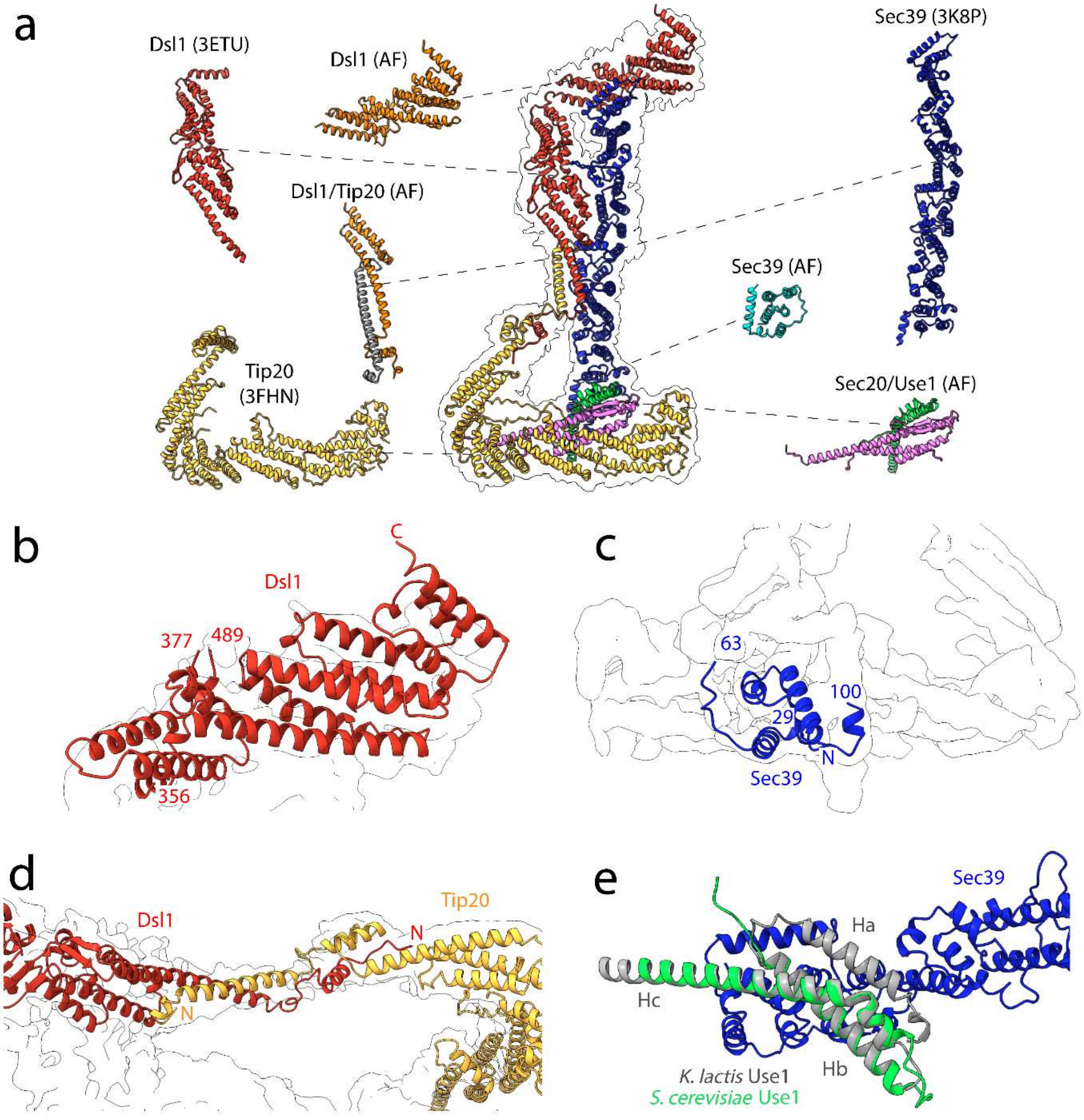


**Extended Data Fig. 5: Crystallographic and Alphafold2-Multimer (AF) contributions to the Dsl1:Qb:Qc complex model.** **a**, Model of the Dsl1:Qb:Qc complex surrounded by an outline of the EM map. Each of the crystal structures (denoted with PDB codes) and AF predictions used in the modeling process are also shown with their relative location specified. **b**, Dsl1ΔN (356-754) was generated by AF to supplement the available crystallographic data. Dsl1ΔN (excluding the lasso 378-488) is overlaid onto the outline of the cryo-EM density of the Dsl1:Qb:Qc complex, to demonstrate fit. **c**, Sec39 (1-100), generated by AF, is overlaid onto the outline of the cryo-EM density of the Dsl1:Qb:Qc complex, to demonstrate fit. **d**, The N-terminal domains of Dsl1 and Tip20 interact directly. The AF-predicted interface of Dsl1 and Tip20 is overlaid onto the outline of the cryo-EM density of the Dsl1:Qb:Qc complex, to demonstrate fit. **e**, AF was used to model *K. lactis* Use1 bound to Sec39, and the resulting prediction was fit into our previously reported electron density map (6WC4). In the resulting model (8FTU), the position of Use1 is the same, but the orientation is flipped, agreeing well with the orientation we observe for *S. cerevisiae* Use1 bound to Sec39.

a

| Protein | pLDDT Scores | Statistics |
|---|---|---|
| Dsl1 334-754 | 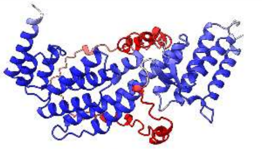 | Average pLDDT: 92.7<br>pTM: 0.715<br>Chimera X Fit CC: 0.9010 |
| Sec39 1-112 | 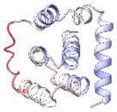 | Average pLDDT: 76.1<br>pTM: 0.657<br>Chimera X Fit CC: 0.8906 |
| Dsl1 1-131,<br>Tip20 1-66 | 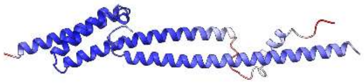 | Average pLDDT: 87.4<br>pTM+ipTM: 0.711<br>Chimera X Fit CC: 0.7363 |
| Sec20 1-184,<br>Use1 2-86 | 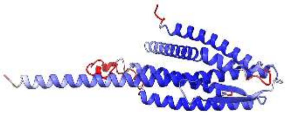 | Average pLDDT: 85.7<br>pTM+ipTM: 0.847<br>Chimera X Fit CC: 0.8584 |

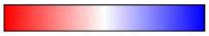


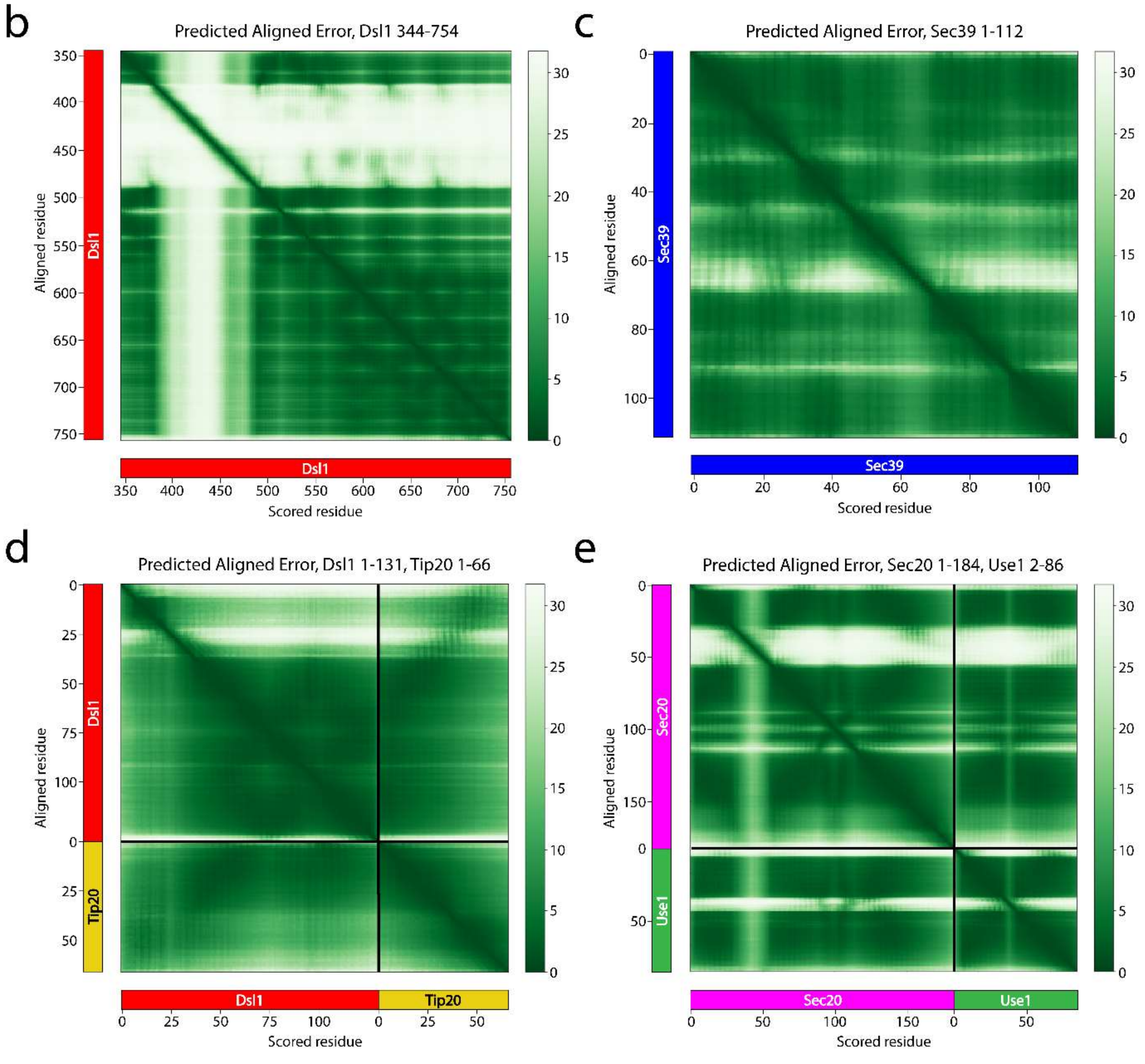

**Extended Data Fig. 6: Statistics on AF contributions to the Dsl1:Qb:Qc complex model. a**, Table listing AF predictions utilized in the model building process of the Dsl1:Qb:Qc complex. *Left:* protein name and residues included in the AF job. *Center*: depiction of the Rank_0 model generated by AF, colored by pLDDT values. Dsl1 (378-488) and Sec20 (38-65), though depicted, were not used in the modeling process as there was no corresponding densities for these regions. *Right*: statistics on the AF jobs. pLDDT was calculated by averaging the score of each residue in a given job used in the final model. pTM (for monomeric jobs) and pTM+ipTM (for multimeric jobs) were extracted from AF directly. CC fit was generated in ChimeraX at the local resolution of the fitted portion of the map. **b**, Predicted aligned error (PAE) graph generated by AF for Dsl1 (344-754). Green indicates a lower distance error for a given residue pair. **c**, PAE graph generated by AF for Sec39 (1-112). **d**, PAE graph generated by AF for Dsl1 (1-131), Tip20 (1-66). **e**, PAE graph generated by AF for Sec20 (1-184), Use1 (2-86).

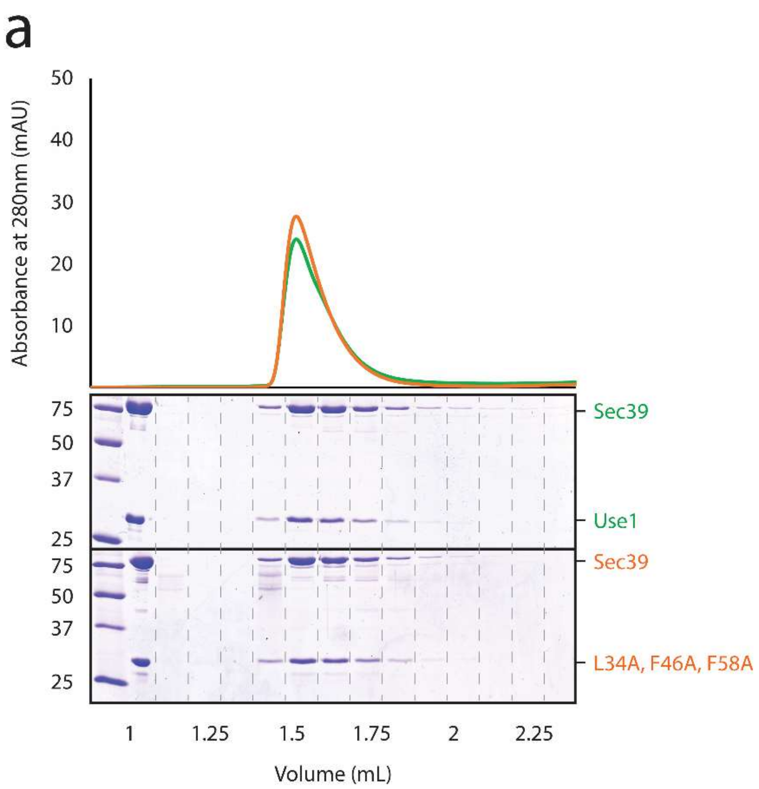


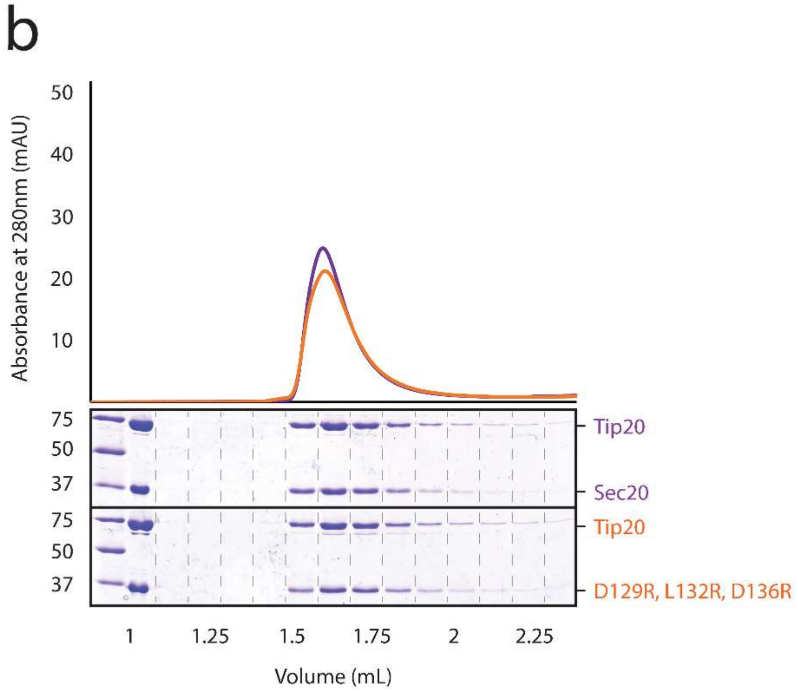


**Extended Data Fig. 7: Mutations in Sec20 and Use1 that abolish SNARE-SNARE binding do not affect protein migration on size-exclusion chromatography. a**, $His_7$-Use1 (1-212):Sec39 migrates with a similar profile to $His_7$-Use1 (1-212) (L34A, F46A, F58A):Sec39 on size-exclusion chromatography. **b**, Sec20-$His_7$ (1-275):Tip20 migrates with a similar profile to Sec20-$His_7$ (1-275) (D129R, L132R, D136R):Tip20.

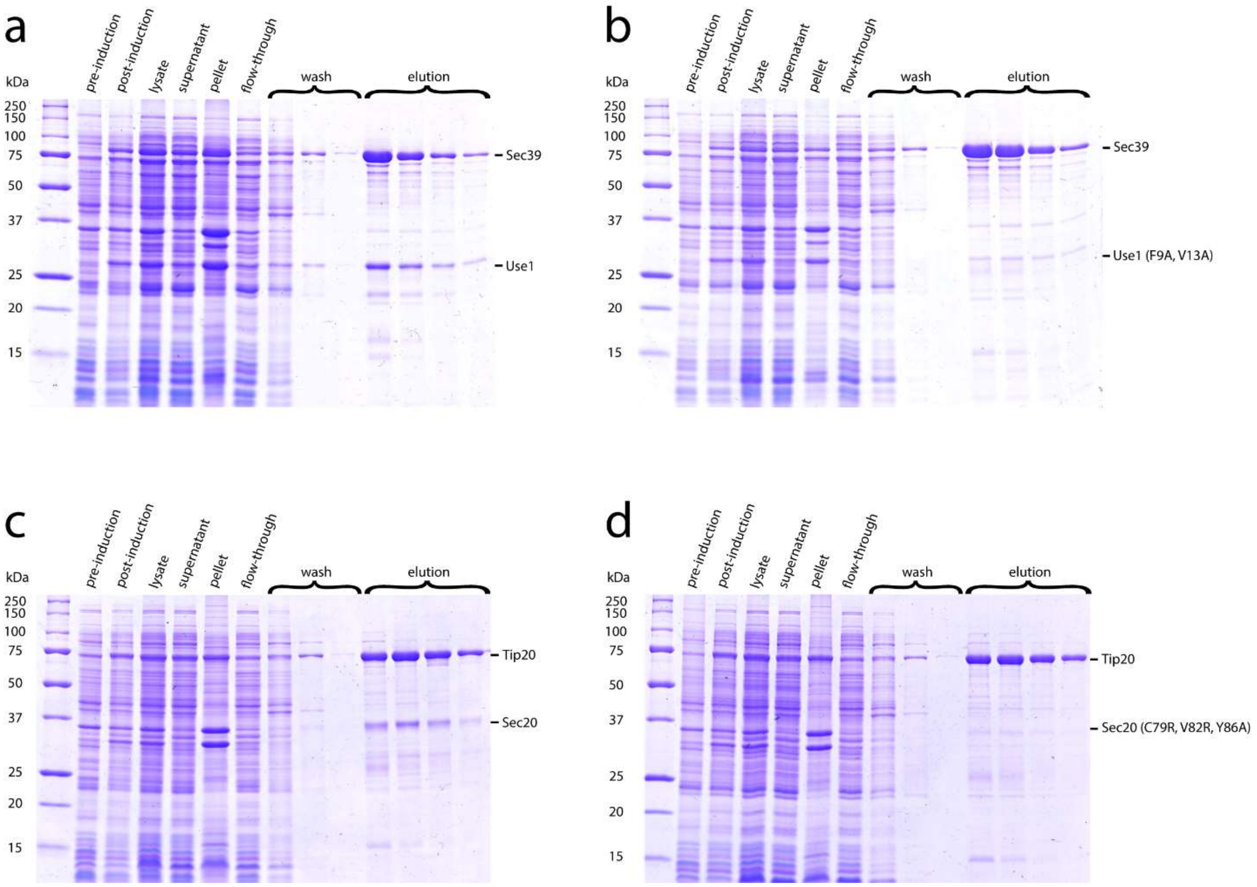


**Extended Data Fig. 8: Mutations designed to disrupt tether:SNARE interactions abolish binding *in vitro*.** **a**, His$_7$-Sec39 pulls down Use1 (1-212). **b**, His$_7$-Sec39 does not pull down mutant Use1 (1-212) (F9A, V13A). **c**, His$_7$-Tip20 pulls down Sec20 (1-275). **d**, His$_7$-Tip20 does not pull down Sec20 (1-275) (C79R, V82R, Y86A).

### Extended Data Table 1. Cryo-EM data collection, refinement, and validation statistics

| | Dsl1:Qb:Qc Complex (EMDB-28204) (PDB 8EKI) |
|---|---|
| **Data collection and processing** | |
| Magnification | 105,000 |
| Voltage (kV) | 300 |
| Electron exposure (e–/Å$^2$) | 45 |
| Defocus range (μm) | -1.25 to -2.5 |
| Pixel size (Å) | 1.114 |
| Symmetry imposed | C1 |
| Initial particle images (no.) | 469,193 |
| Final particle images (no.) | 49,947 |
| Map resolution (Å) | 4.5 |
| FSC threshold | 0.143 |
| Map resolution range (Å) | 3.9 to 10.7 |
| **Refinement** | |
| Initial model used (PDB code) | 3ETU, 3K8P, 3FHN, AlphaFold2 |
| Model resolution (Å) | 4.5 |
| FSC threshold | 0.143 |
| Model resolution range (Å) | 3.9 to 10.7 |
| Map sharpening *B* factor (Å$^2$) | -189.2 (Mask 1, EMD-29447)<br>-477.6 (Mask 2, EMD-28760)<br>-316.2 (Mask 3, EMD-28768)<br>-375.3 (Mask 4, EMD-28762) |
| Model composition | |
| Non-hydrogen atoms | 18750 |
| Protein residues | 2278 |
| Ligands | - |
| *B* factors (Å$^2$) | |
| Protein | 339.24 |
| Ligand | - |
| R.m.s. deviations | |
| Bond lengths (Å) | 0.005 |
| Bond angles (°) | 0.740 |
| Validation | |
| MolProbity score | 2.34 |
| Clashscore | 27.24 |
| Poor rotamers (%) | 0.56 |
| Ramachandran plot | |
| Favored (%) | 93.74 |
| Allowed (%) | 6.13 |
| Disallowed (%) | 0.13 |
